# Royal decree: gene expression in transgenerationally immune primed bumblebee workers mimics a primary immune response

**DOI:** 10.1101/043638

**Authors:** Seth M. Barribeau, Paul Schmid-Hempel, Ben M. Sadd

## Abstract

Invertebrates lack the cellular and physiological machinery of the adaptive immune system, but show specificity in their immune response [1, 2] and immune priming [3-11]. Functionally, immune priming is comparable to immune memory in vertebrates. Individuals that have survived exposure to a given parasite are better protected against subsequent exposures. Protection may be cross-reactive (e.g. [12]), but demonstrations of persistent and specific protection in invertebrates are increasing [3, 5]. This immune priming can cross generations ("trans-generational" immune priming) [4, 8], preparing offspring for the prevailing parasite environment. While these phenomena gain increasing support, the mechanistic foundations underlying such immune priming, both within and across generations, remain largely unknown. Using a transcriptomic approach, we show a bacterial challenge to bumblebee queens, known to induce trans-generational immune priming, alters daughter (worker) gene expression. Daughters, even when unchallenged themselves, constitutively express a core set of the genes induced upon direct bacterial exposure, including high expression of antimicrobial peptides, a beta-glucan receptor protein implicated in bacterial recognition and the induction of the *toll* signaling pathway[13], and *slit-3* which is important in honeybee immunity[14]. Maternal challenge results in a distinct upregulation of their daughters’ immune system, with a signature overlapping with the induced individual response to a direct immune challenge. This will mediate mother-offspring protection, but also associated costs related to reconfiguration of constitutive immune expression. Identification of conserved immune pathways in memory-like responses has important implications for our understanding of the innate immune system, including the innate components in vertebrates, which share many of these pathways[15].

**Author Summary:** Invertebrate individuals surviving exposure to an infectious disease can become better at fighting future infection by that same disease. This protection, known as immune priming, can even be transferred to the individuals’ offspring. The functional outcome is very similar to that of vertebrate immune memory, but the mechanisms of how invertebrates achieve immune priming within an individual or across generations remain enigmatic. We found that bumblebee daughters of mothers exposed to a simulated bacterial infection express strongly many of the genes that they would activate if they were themselves infected. Our results show how immune priming across generations might be produced in bumblebees. Many parts of the invertebrate immune system are shared with us, and thus our study also sheds a light on how diverse immune memory-like effects could be achieved.

## Introduction

Parasites, broadly construed to include both macro-and microparasites, are ubiquitous and can cause significant damage to their hosts. As a consequence, parasites represent a major selective force for any organism. Hosts, in turn, have adaptations that prevent parasite establishment and reduce the costs of having an infection. These adaptations, which can be broadly viewed as elements of a defense system, notably including the immune response, range in their specificity, their mode of action, and the nature of regulation. As investment into immunity is costly on multiple levels[16], the most efficient investment into immunity will be a function of the prevailing pressure from parasites (likelihood of encounter and virulence) and demands imposed by other life-history traits. On an ecological scale, there will therefore be a benefit to a plastic adjustment of immune investment relevant for sufficiently accurate "perceived" risk of parasitism. This perception may be related to ecological conditions, such as crowding[17], but may also result from prior immunological experience with parasites. In particular, hosts can encounter the same parasites multiple times within their lifetime, and across generations. If hosts encounter the same parasite repeatedly, some form of memory, which would improve resistance to that same parasite upon reexposure, will be adaptive.

The best-studied and classic example of an adjustment in immune responses in relation to a prior parasite exposure is the adaptive immune system of vertebrates. The adaptive immune system, which produces specific and long-lasting protection against subsequent exposure to the same parasite, is based on a repertoire of specialized lymphocytes and its molecular underpinnings are well characterized[18]. There is growing evidence that functionally comparable processes may exist in other organisms[19, 20]. To avoid mechanism-based confusion in terms, these phenomena so far described for several invertebrates[19], plants[20, 21] or even bacteria[22], are frequently referred to as 'immune priming'. Astonishingly, induced protection against parasites in these systems can traverse generations, a phenomenon known as trans-generational immune priming [23-25].

The molecular understanding of immune priming outside of the adaptive immune system of vertebrates is still in its infancy. Some progress has been made in understanding these mechanisms insects[26, 27] and plants[28]. Invertebrates are particularly important to understand in this regard as they share a number of conserved characteristics of the innate immune system with vertebrates, including humans[29]. The potential for these innate immune components to exhibit a memory-like response is an intriguing possibility[30, 31]. While invertebrates may serve as a valuable model for understanding memory-like phenomena produced solely by innate immune system, the mechanisms remain enigmatic. Studies have identified the roll of the *toll* pathway and phagocytosis within an individual’s life[10] in *Drosophila melanogaster;* and cellular mechanisms are suspected for mosquitoes [9].

Here we investigate patterns of gene expression underlying the phenomenon of trans-generational immune priming in a social insect, the bumblebee *Bombus terrestris*. In social insects, such as bumblebees, temporal and spatial overlap of worker offspring and their mothers will mean that they are faced with a parasite threat that can, with a high probability, be predicted from the mother’s prior immunological experience. *B*. *terrestris*, is a model of ecological host-parasite interactions that shows a specific immune response [1, 2, 32, 33], and within-individual [5, 34] and trans- generational [4, 6] immune priming. Daughters of bacterial-challenged queens show elevated antibacterial responses, but pay costs in terms of resistance to distinct parasites[5, 7]. The mechanisms underlying these responses are unknown.

We injected *B. terrestris* queens with a heat-inactivated inoculum of the Gram-positive bacterium *Arthobacter globiformis*, in the same manner as previous studies demonstrating trans-generational immunity[6, 7], To gain some insight into the molecular foundations of observed trans-generational immunity in this system we measured genome-wide expression of subsequently produced naïve daughters (*Arthrobacter-Naïve* [AN] treatment) relative to the expression of naïve daughters born from unchallenged mothers (Naïve-Naïve [NN] treatment). We further contrast this memory response with the immune response of daughters that are exposed to the bacterial challenge, but whose mothers were naïve (*Naïve-Arthrobacter* [NA] treatment).

## Results

Whole genome expression, as measured by mRNA sequencing on the Illumina HiSeq platform revealed that when workers from unchallenged, naïve, mothers were directly challenged with the bacterial inoculum (NA) they responded with significant differential expression of 327 genes (Table S1). Naïve workers from challenged mothers (AN) significantly altered the expression of only 21 genes (Table S2), but 20 of these are shared with the direct induced response (NA) (Fig 1). These shared genes (Fig 2) include all known bumblebee antimicrobial peptides (*abaecin*, two *apidaecins*, *defensin*, *hymenoptaecin*) and a number of additional known immune genes such as *battenin*, *laccase-2*, *slit-3*, and *yellow*. The only gene differentially expressed in the primed condition, but not under direct induction, codes for LOC100644816, a 53aa hydrophobic (58.49% of residues) peptide with homology to Mast Cell Degranulating Peptide (MCDP) from another bumblebee, *B. pennsylvanicus*.

**Figure 1:**
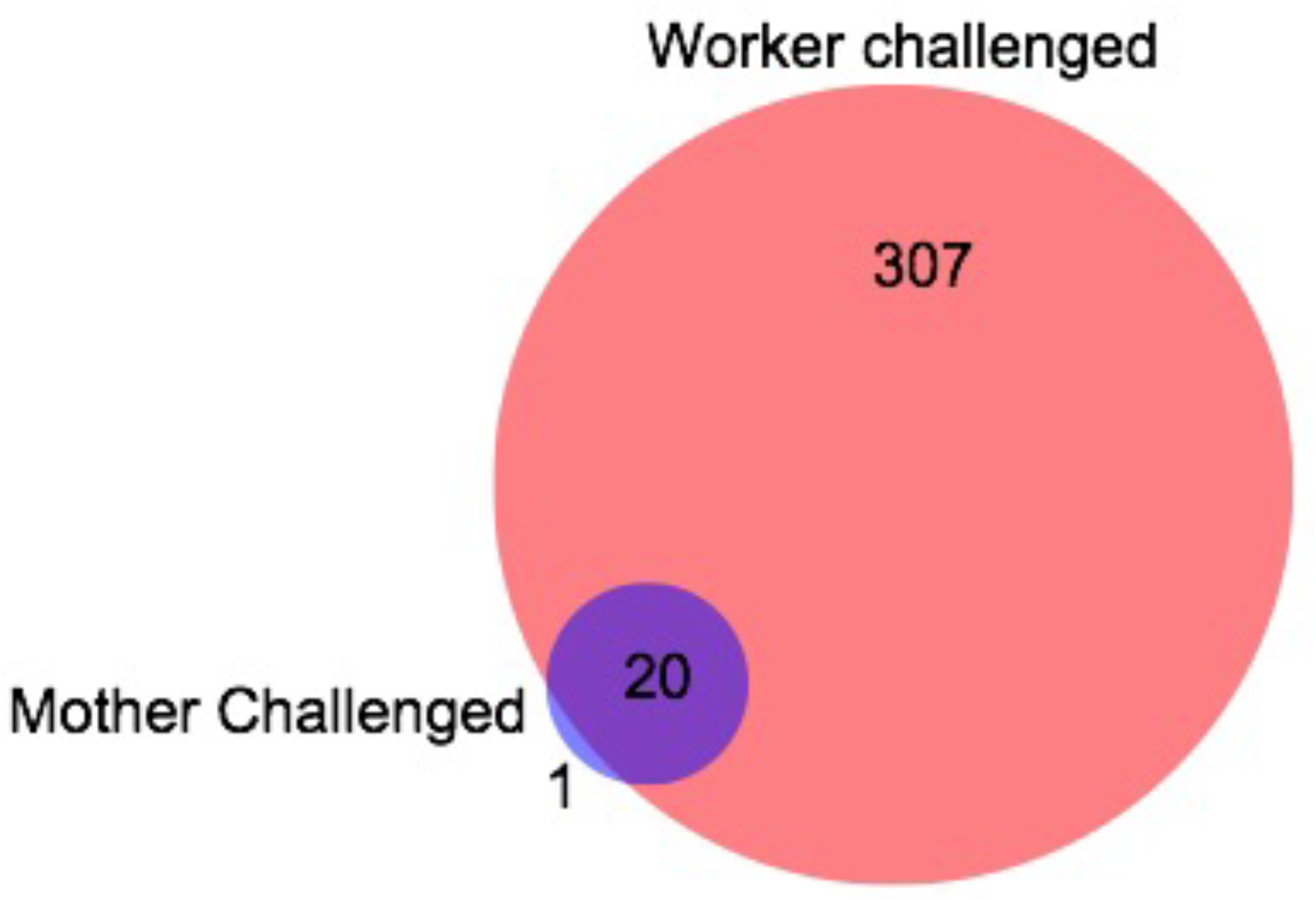
The number of differentially expressed genes in naïve worker offspring of mother queens that were injected with heat killed Gram-positive bacterium (*Arthrobacter globiformis*) (trans-generational immunity treatment; AN), and worker offspring from naïve mother queens but themselves exposed to an immune challenge of *A. globiformis* (induced immune response condition; NA). The expression of these genes is measured relative to that of naïve worker offspring of naïve mothers (NN).

**Figure 2:**
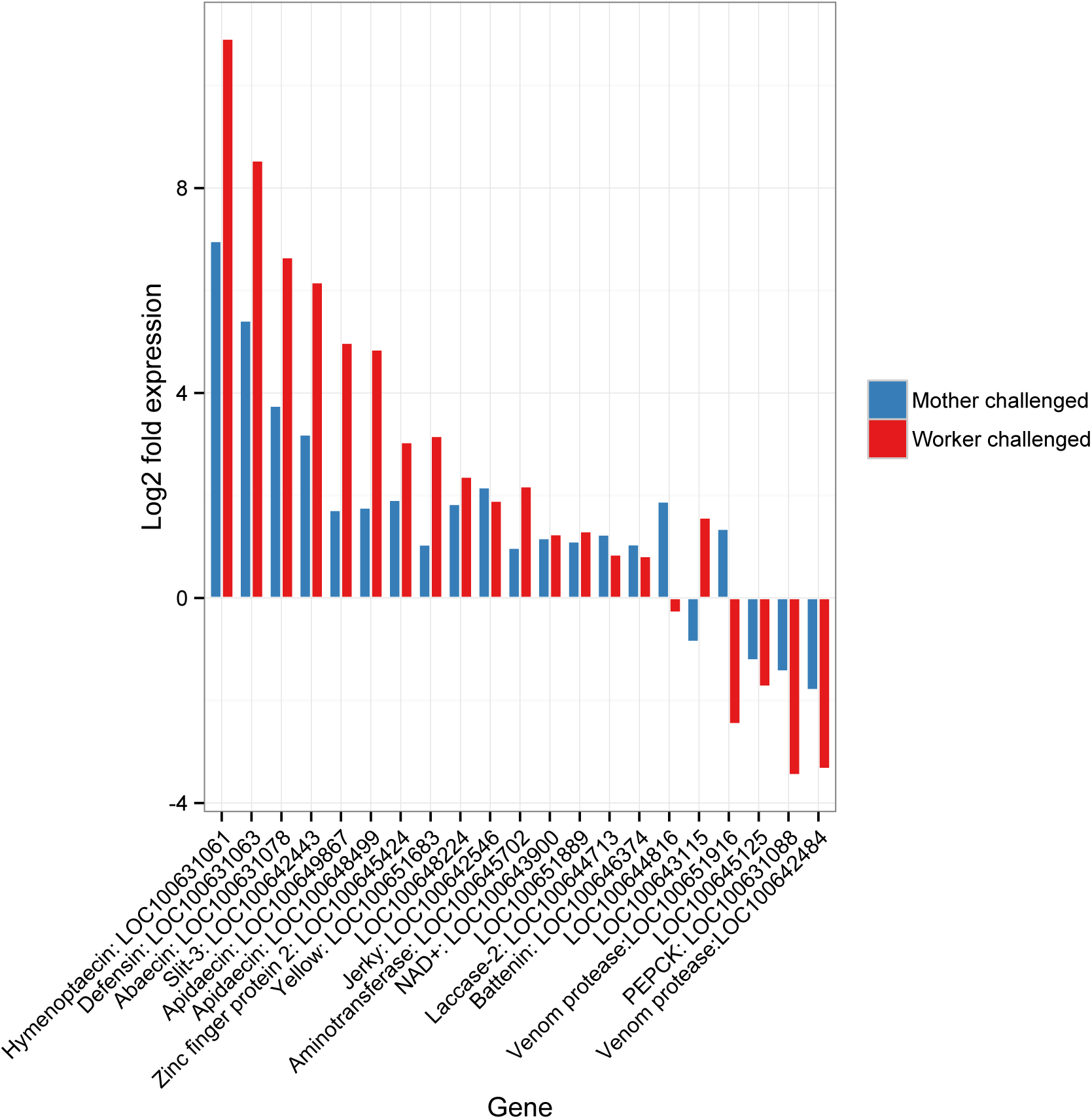
Log 2 fold expression based on RNAseq data (relative to naïve worker offspring from naïve mother queens) for all genes that are significantly differentially expressed in the trans-generational priming condition (naïve offspring of challenged mothers, blue). We also show the expression of challenged workers from naïve mothers (red) to demonstrate the similarity of the induced response to a direct challenge to the signature of trans-generational immunity. All differentially expressed genes here are also significantly differentially expressed upon direct challenge, except for LOC100644816, which encodes for mast cell degranulating peptide. qPCR confirmation of these results can be found in Fig. S1.

We confirmed the patterns determined by the whole genome transcriptome approach (Fig 2) by targeted qPCR of a suite of immune genes (Table S3). Our qPCR results agree with our transcriptomic results for all tested genes. This included the high constitutive expression of the antimicrobial peptides and additionally of a beta-glucan receptor protein (BGRP, Fig S1, Table S4) in naïve offspring of immune-challenged mothers (AN). There was a trend for higher BGRP expression in the transcriptome of workers from challenged mothers, but this was not significant after correction for multiple testing (*P* < 0.01 before correction, 0.072 after correction). Differential expression of this receptor may be particularly relevant as it can trigger the *toll* signaling pathway and downstream antimicrobial peptide production.

We identified a number of different isoforms for putative immune response genes, including for antimicrobial peptides (Fig S2-17: *abaecin* [LOC100631078], 4 isoforms; both *apidaecins* [LOC100649867], 2 isoforms, and [LOC100648499], 3 isoforms, aminopeptidase [LOC100645702], tetraspannin [LOC100651747], a venom protease [LOC100651916], an uncharacterized protein shared only within honeybees and bumblebees [LOC100645125], and a novel gene [NC_015763.1:3848320-3855802] with sequence homology to *A. mellifera* cuticular protein 14. We also identified two *dscam* like genes with multiple isoforms (Supplemental Fig S18-21; LOC100644003, 12 isoforms; LOC100649765, 9 isoforms). Among the significantly differentially expressed genes, isoform transcript abundance did not vary significantly among conditions.

## Discussion

We found that offspring workers which had never been exposed themselves ("naive workers") but whose mothers were exposed to a bacterial immune challenge express a strikingly pathogen exposure-like immune response, as compared to offspring workers from naïve mothers. In fact, all but one of the differentially expressed genes in this priming condition were shared with workers that were directly immune challenged with the same bacterium. This indicates a major reconfiguration of the constitutively expressed immune gene profile, and is one that will likely confer appropriate benefits in the face of specific parasites, but may also result in the costs previously described when there is mismatch between the maternal parasite environment and the offspring parasite environment[7]. These results give us an insight into the innate immune-related molecular pathways at the base of invertebrate immune priming across generations.

Among the differentially expressed genes, all of the antimicrobial peptides are upregulated. It is particularly noteworthy that these are the end-point of the immune response, indicating an immediate readiness of defense in trans-generationally primed individuals. We also find increased expression in a number of other immunologically important genes including *yellow* and *laccase-2*, which are involved in the melanization response [35, 36], *battenin*, the *D. melanogaster* homolog of which (*CLN3*) regulates *JNK* signaling[37], and *slit-3* which is induced upon bacterial challenge in honeybees and the leaf-cutting ant *Atta cephalotes* (a.k.a. *IRP30*)[14]. We also found that a beta-glucan receptor protein (BGRP) was more highly expressed in naïve workers of challenged queens. BGRPs induce the *toll* signaling pathway in invertebrates[13]. The only gene that was differentially expressed in the primed condition, but not in directly challenge workers, was the mast cell degranulating peptide (MCDP), which is found in venom[38]. MCDP, neuro-and immunotoxic, is named for its degranulating effect on vertebrate granulocytes[38, 39]. Whether this peptide also has the same effect on invertebrate granulocytes (a class of haemocytes) that are important for phagocytosis[40] is unknown. Intra-generationally primed *Drosophila melanogaster*, utilize the *toll* pathway and phagocytosis, but not antimicrobial peptides[10] that appear to play an important role here.

The down syndrome cell adhesion molecule (*dscam*) is implicated in immune defenses, and because of its ability to produce prodigious numbers of isoforms[25] has been proposed as a possible mechanism for specific immune memory[41]. We detect two *dscam* like genes that produce multiple isoforms. These genes nor their isoforms are not, however, differentially expressed in the priming condition or in workers that are directly exposed to the bacterium. While this does not rule out a role for *dscam* isoforms in immune priming, it suggests that differential expression of isoforms is not a major component of trans-generational antibacterial immune priming in this system.

The elevated constitutive gene expression into adulthood of a holometabolous insect, with its associated tissue rearrangements, is testament to the persistence of the trans-generational priming in the innate immune system. Evidence in insects of elevated constitutive expression of immune-related genes that is precipitated by immune experiences in prior generations is important beyond a demonstration of the underlying mechanistic foundations of trans-generational immunity. It will also have important consequences for the fitness costs associated with this phenomenon, which will influence the conditions under which it may be expected to evolve and be maintained by selection. Elevated immune investment comes at a cost to an organism through resource trade-offs and other routes [42]. Higher constitutive expression of immunity in naïve offspring may constrain their investment into other life-history traits, especially under conditions where resources are scarce. The striking signature of gene expression related to trans-generational immunity is also likely to underpin other related costs, including increased susceptibility to a distinct parasite infection, as has been demonstrated in this system [7].

Evidence is mounting that the evolutionarily ancient innate immune system is able to retain information about immune history in both vertebrates and invertebrates[43], which translates to better defenses upon subsequent exposure. This priming effect is observed both within the lifetime of an individual and between parents and offspring. Trans-generational immune priming likely evolved as a part of parental care and investment into offspring. This may be particularly important in social insects, such as *B. terrestris*, where generations overlap and related individuals very intimately share an environment - including parasites - in a closed, populous, highly interactive colony. While our study does not attempt to identify the mechanisms involved in transfer of immune compounds to the offspring, a recent paper in honeybees identified the yolk protein vitellogenin as playing a role in binding and transferring bacterial cell components to eggs[44]. Here we find that trans-generationally primed workers - even if not infected themselves - increase transcription of antimicrobial peptides (that in part are under the control of the *toll* signaling pathway) and a key recognition protein that induces *toll* signaling. This transcriptional signature resembles an abridged version of the normal response to infection, suggesting that *B. terrestris* achieves trans-generational protection by sequestering the existing induced responses into prophylactic constitutive expression to prevent parasite establishment. A recent study in moths found elevated ovary expression of some immune genes in daughters of challenged mothers, hinting that these responses may even be transmitted across multiple generations[45]. The conserved nature of these innate immune pathways suggests that the patterns detected here may also underlie immune experience based immune system plasticity not only invertebrates, but also in the innate immune system of vertebrates.

## Materials and Methods

### Experimental methods

We collected queens as they emerged from hibernation in spring 2013 in northern Switzerland and maintained them under standard colony establishment conditions[1]. All of the colonies used for this experiment were microscopically checked for common infections twice and found to be clear of identifiable infection. On their production by the colonies, young queens (gynes) were removed and mated to males from unrelated colonies. We designed the matings such that sister queens from one colony were mated to males all derived from a single colony to produce comparable genetic backgrounds for matching across treatments. Five days after mating, we hibernated the queens for 48 days at 4C. Seven-days after removal from hibernation we either injected queens to the challenged with 2 μl of 10^8^ colony-forming-units/mL of *Arthrobacter globiformis* (DSM 20124) that had been heat inactivated by heating at 95C for 5 min, washed three times and resuspended in ringer saline solution. Naive, unchallenged queens were handled similarly but not injected. We then allowed queens to found colonies in the lab. We used three sister queen colony sets. We deliberately used these second generation colonies to exclude unknown maternal effects outside of our treatments. Emerging adult worker offspring from naïve queens were uniformly distributed to a naive group (NN) or an induced treatment (NA). In the induced treatment, daughters five-days post-eclosion received an injection of 2 μl of 10^8^ colony-forming-units/mL of *A. globiformis* prepared as above replicated twice per condition, per colony, (NA: naïve queens, *A. globiformis* exposed worker daughters) and were snap frozen in liquid nitrogen 24hrs after injection. Naïve group workers (NN) were handled similarly, but not injected, and frozen at the same time. Similarly, we took workers from queens that were exposed to the bacterial challenge and handled and froze them as above (AN). We extracted RNA from the workers following the same protocols as in [1] but using whole abdomens. For RNA sequencing we pooled the RNA from two individual workers per queen and treatment combination. The sequencing used the Illumina HiSeq 2000 platform and was done at the Beijing Genomics Institute.

After removing adapters and poor quality reads we mapped the reads to the *B. terrestris* genome[46] with Tophat2[47] in two ways. First, using the annotated transcripts (-G option) to assess differential expression of known genes, and second, without this restriction to assess isoform variation. We identified differentially expressed genes using Cuffdiff[48, 49]. In both cases we used the current version of the *B. terrestris* genome (Bterr_1.0) with the accompanying gtf annotation file. We limited the maximum intron size to 50kb. The analyses compared the expression of naïve daughters of challenged mothers (AN) vs naïve daughters of naïve mothers (NN), and separately compared the induced response (NA) to the baseline expression of NN workers.

From un-pooled offspring samples described above, we also synthesized cDNA using the QuantiTect Reverse Transcription Kit (Qiagen) following the manufacturers instructions. In addition, cDNA was synthesized from offspring of queens from three additional matched genetic background providing further AN, NA and NN samples. One of these colonies had duplicate offspring for each treatment, while the other two colonies had one replicate offspring. Prior to cDNA synthesis potential DNA contamination was removed from all RNA samples using the Turbo DNA-free kit (Ambion) according to the manufacturers instructions. In the reverse transcribed samples, we quantified the expression of 25 immune genes relative to two invariant housekeeping genes (elongation factor 1α and ribosomal protein L13 based on their scores in geNorm, qBase plus, biogazelle) and analyzed as in[1]. Full details of these genes and their primers are in Table S5. We used the mean difference in expression of the target gene from the composite housekeeping gene (dCt) from each colony for subsequent analyses. We transformed the mean dCt value for each gene using Yeo-Johnson transformations to improve normality and homoscedasticity and used paired t-tests within colony genetic background to assess statistical differences between NN and AN treatments and between NN and NA treatments (Table S6).

## Acknowledgements

We thank Miguel Jales and Elke Karaus for their technical assistance, and Nicole Gerardo for her comments on an earlier draft of this manuscript. Some of these data were generated at the Genetic Diversity Centre of ETH Zürich. We thank the Bumblebee Genome Consortium (http://hymenopteragenome.org/beebase/) for providing genomic resources that were used for this study.

## References

1. Barribeau SM, Sadd BM, du Plessis L, Schmid-Hempel P. Gene expression differences underlying genotype-by-genotype specificity in a host-parasite system. P Natl Acad Sci USA. 2014;111(9):3496-501.

2. Riddell CE, Adams S, Schmid-Hempel P, Mallon EB. Differential expression of immune defences is associated with specific host-parasite interactions in insects. PLoS One. 2009;4 (10). doi: ARTNe7621DOI10.1371/journal.pone.0007621. PubMed PMID: ISI:000271147400017.

3. Roth O, Sadd BM, Schmid-Hempel P, Kurtz J. Strain-specific priming of resistance in the red flour beetle, Tribolium castaneum. Proc R Soc Lond B. 2009;276(1654):145-51. doi: Doi 10.1098/Rspb.2008.1157. PubMed PMID: ISI:000262004900019.

4. Sadd BM, Kleinlogel Y, Schmid-Hempel R, Schmid-Hempel P. Trans-generational immune priming in a social insect. Biol Lett. 2005;1(4):386-8. Epub 2006/12/07. doi: H762604181757515[pii]10.1098/rsbl.2005.0369. PubMed PMID: 17148213; PubMed Central PMCID: PMC1626361.

5. Sadd BM, Schmid-Hempel P. Insect immunity shows specificity in protection upon secondary pathogen exposure. Curr Biol. 2006;16(12):1206-10.

6. Sadd BM, Schmid-Hempel P. Facultative but persistent transgenerational immunity via the mother's eggs in bumblebees. Curr Biol. 2007;17 (24):R1046-R7. doi: Doi 10.1016/J.Cub.2007.11.007. PubMed PMID: WOS:000251852200012.

7. Sadd BM, Schmid-Hempel P. A distinct infection cost associated with trans-generational priming of antibacterial immunity in bumble-bees. Biol Lett. 2009;5 (6):798-801.

8. Roth O, Joop G, Eggert H, Hilbert J, Daniel J, Schmid-Hempel P, et al. Paternally derived immune priming for offspring in the red flour beetle, Tribolium castaneum. J Anim Ecol. 2010;79(2):403-13. doi: 10.1111/j.1365-2656.2009.01617.x. PubMed PMID: 19840170.

9. Rodrigues J, Brayner FA, Alves LC, Dixit R, Barillas-Mury C. Hemocyte differentiation mediates innate immune memory in Anopheles gambiae mosquitoes. Science. 2010;329 (5997):1353-5. doi: 10.1126/science.1190689. PubMed PMID: 20829487; PubMed Central PMCID: PMC3510677.

10. Pham LN, Dionne MS, Shirasu-Hiza M, Schneider DS. A specific primed immune response in Drosophila is dependent on phagocytes. PLoS Pathog. 2007;3(3):e26. doi: 10.1371/journal.ppat.0030026. PubMed PMID: 17352533; PubMed Central PMCID: PMC1817657.

11. Boman HG, Nilsson I, Rasmuson B. Inducible antibacterial defence system in Drosophila. Nature. 1972;237 (5352):232-5. PubMed PMID: 4625204.

12. Moret Y, Siva-Jothy MT. Adaptive innate immunity? Responsive-mode prophylaxis in the mealworm beetle, Tenebrio molitor. P Roy Soc Lond B Bio. 2003;270 (1532):2475-80. doi: Doi 10.1098/Rspb.2003.2511. PubMed PMID: ISI:000187086200009.

13. Gottar M, Gobert V, Matskevich AA, Reichhart JM, Wang C, Butt TM, et al. Dual detection of fungal infections in Drosophila via recognition of glucans and sensing of virulence factors. Cell. 2006;127 (7):1425-37. doi: 10.1016/j.cell.2006.10.046. PubMed PMID: 17190605; PubMed Central PMCID: PMC1865096.

14. Albert S, Gatschenberger H, Azzami K, Gimple O, Grimmer G, Sumner S, et al. Evidence of a novel immune responsive protein in the Hymenoptera. Insect Biochem Mol Biol. 2011;41 (12):968-81. doi: 10.1016/j.ibmb.2011.09.006. PubMed PMID: 22001069.

15. Silverman N, Maniatis T. NF-kappaB signaling pathways in mammalian and insect innate immunity. Genes Dev. 2001;15 (18):2321-42. doi: 10.1101/gad.909001. PubMed PMID: 11562344.

16. Schmid-Hempel P. Variation in immune defence as a question of evolutionary ecology. P R Soc B. 2003;270 (1513):357-66. doi: papers2://publication/uuid/831D6BD2-BFFA-4C1B-AD05-D0958A128B1B.

17. Wilson K, Thomas MB, Blanford S, Doggett M, Simpson SJ, Moore SL. Coping with crowds: density-dependent disease resistance in desert locusts. P Natl Acad Sci USA. 2002;99 (8):5471-5. doi: papers2://publication/doi/10.1073/pnas.082461999.

18. Janeway C, Travers P, Walport M, Shlomchik M. Immunobiology. New York: Garland Science; 2001.

19. Kurtz J. Specific memory within innate immune systems. Trends Immunol. 2005;26(4):186-92. doi: 10.1016/j.it.2005.02.001. PubMed PMID: 15797508.

20. Pastor V, Luna E, Mauch-Mani B, Ton J, Flors V. Primed plants do not forget. Environ Exp Bot. 2013;94:46-56.

21. Mou Z, Fan W, Dong X. Inducers of plant systemic acquired resistance regulate NPR1 function through redox changes. Cell. 2003;113 (7):935-44. doi: papers2://publication/uuid/11A9A2F9-06C6-4D86-93A1-D245D003515F.

22. Horvath P, Barrangou R. CRISPR/Cas, the immune system of bacteria and archaea. Science. 2010;327 (5962):167-70. doi: 10.1126/science.1179555. PubMed PMID: 20056882.

23. Kleino A, Valanne S, Ulvila J, Kallio J, Myllymaki H, Enwald H, et al. Inhibitor of apoptosis 2 and TAK1-binding protein are components of the Drosophila Imd pathway. EMBO J. 2005;24(19):3423-34. doi: Doi 10.1038/Sj.Emboj.7600807. PubMed PMID: W0S:000232551800008.

24. Nakamoto M, Moy RH, Xu J, Bambina S, Yasunaga A, Shelly SS, et al. Virus Recognition by Toll-7 Activates Antiviral Autophagy in Drosophila. Immunity. 2012;36(4):658-67. doi: Doi 10.1016/J.Immuni.2012.03.003. PubMed PMID: WOS:000303223900017.

25. Yu HH, Yang JS, Wang J, Huang Y, Lee T. Endodoamin Diversity in the Drosophila Dscam and Its Roles in Neuronal Morphogenesis. J Neurosci. 2009;29(6):1904-14. doi: Doi 10.1523/Jneurosci.5743-08.2009. PubMed PMID: WOS:000263301200032.

26. Christofi T, Apidianakis Y. Drosophila immune priming against Pseudomonas aeruginosa is short-lasting and depends on cellular and humoral immunity. F1000Res 2013;2:76. doi: 10.12688/f1000research.2-76.v1. PubMed PMID: 24358857; PubMed Central PMCID: PMC3752738.

27. Pham LN, Dionne MS, Shirasu-Hiza M, Schneider DS. A Specific Primed Immune Response in Drosophila Is Dependent on Phagocytes. PLoS Path. 2007;3 (3):e26. doi: papers2://publication/doi/10.1371/journal.ppat.0030026.

28. Muthamilarasan M, Prasad M. Plant innate immunity: an updated insight into defense mechanism. J Biosci. 2013;38(2):433-49. PubMed PMID: 23660678.

29. Lemaitre B. The road to Toll. Nature Reviews Immunology. 2004. doi: papers2://publication/uuid/2902462C-CA07-4841-AB6C-90646B275BC1.

30. Sun JC, Ugolini S, Vivier E. Immunological memory within the innate immune system. The EMBO journal. 2014;33 (12):1295-303. doi: 10.1002/embj.201387651. PubMed PMID: 24674969; PubMed Central PMCID: PMC4194120.

31. Chambers MC, Schneider DS. Pioneering immunology: insect style. Curr Opin Immunol. 2012;24 (1):10-4. doi: papers2://publication/doi/10.1016/j.coi.2011.11.003.

32. Barribeau SM, Schmid-Hempel P. Qualitatively different immune response of the bumblebee host, Bombus terrestris, to infection by different genotypes of the trypanosome gut parasite, Crithidia bombi. Infect, Genet Evol. 2013;20:249-56.

33. Sadd BM, Barribeau SM. Heterogeneity in infection outcome: lessons from a bumblebee-trypanosome system. Parasite Immunol. 2013. doi: 10.1111/pim.12043. PubMed PMID: 23758554.

34. Moret Y, Schmid-Hempel P. Immune defence in bumble-bee offspring. Nature. 2001;414(6863):506. doi: 10.1038/35107138. PubMed PMID: 11734840.

35. Ferguson LC, Green J, Surridge A, Jiggins CD. Evolution of the insect yellow gene family. Mol Biol Evol. 2011;28(1):257-72. doi: 10.1093/molbev/msq192. PubMed PMID: 20656794.

36. Arakane Y, Muthukrishnan S, Beeman RW, Kanost MR, Kramer KJ. Laccase 2 is the phenoloxidase gene required for beetle cuticle tanning. Proc Natl Acad Sci U S A. 2005;102 (32):11337-42. doi: 10.1073/pnas.0504982102. PubMed PMID: 16076951; PubMed Central PMCID: PMC1183588.

37. Tuxworth RI, Vivancos V, O'Hare MB, Tear G. Interactions between the juvenile Batten disease gene, CLN3, and the Notch and JNK signalling pathways. Hum Mol Genet. 2009;18(4):667-78. doi: 10.1093/hmg/ddn396. PubMed PMID: 19028667; PubMed Central PMCID: PMC2638826.

38. Argiolas A, Herring P, Pisano JJ. Amino acid sequence of bumblebee MCD peptide: a new mast cell degranulating peptide from the venom of the bumblebee Megabombus pennsylvanicus. Peptides. 1985;6 Suppl 3:431-6. PubMed PMID: 2421265.

39. Ziai MR, Russek S, Wang HC, Beer B, Blume AJ. Mast cell degranulating peptide: a multi-functional neurotoxin. The Journal of pharmacy and pharmacology. 1990;42 (7):457-61. PubMed PMID: 1703229.

40. Strand MR. Insect Hemocytes and Their Role in Immunity. In: Beckage NE, editor. Insect Immunology. San Diego: Elsevier; 2008. p. 25-47.

41. Armitage SAO, Peuβ R, Kurtz J. Dscam and pancrustacean immune memory - A review of the evidence. Dev Comp Immunol. 2014;in press.

42. Sadd BM, Schmid-Hempel P. Principles of ecological immunology. Evol Appl. 2009;2 (1):113-21. doi: Doi 10.1111/J.1752-4571.2008.00057.X. PubMed PMID: ISI:000262827800012.

43. Netea MG, Quintin J, van der Meer JW. Trained immunity: a memory for innate host defense. Cell host & microbe. 2011;9 (5):355-61. doi: 10.1016/j.chom.2011.04.006. PubMed PMID: 21575907.

44. Salmela H, Amdam GV, Freitak D. Transfer of Immunity from Mother to Offspring Is Mediated via Egg-Yolk Protein Vitellogenin. PLoS Path. 2015. Epub July 31. doi: 10.1371/journal.ppat.1005015.

45. Trauer-Kizilelma U, Hilker M. Insect parents improve the anti-parasitic and anti-bacterial defence of their offspring by priming the expression of immune-relevant genes. Insect Biochem Mol Biol. 2015;64:91-9. doi: 10.1016/j.ibmb.2015.08.003. PubMed PMID: 26255689.

46. Sadd BM, Barribeau SM, Bloch G, de Graaf DC, Dearden P, Elsik CG, et al. The genomes of two key bumblebee species with primitive eusocial organization. Genome Biol. 2015;16:76. doi: 10.1186/s3059-015-0623-3. PubMed PMID: 25908251; PubMed Central PMCID: PMC4414376.

47. Kim D, Pertea G, Trapnell C, Pimentel H, Kelley R, Salzberg SL. TopHat2: accurate alignment of transcriptomes in the presence of insertions, deletions and gene fusions. Genome Biol. 2013;14(4):R36. doi: 10.1186/gb-2013-14-4-r36. PubMed PMID: 23618408.

48. Trapnell C, Williams BA, Pertea G, Mortazavi A, Kwan G, van Baren MJ, et al. Transcript assembly and quantification by RNA-Seq reveals unannotated transcripts and isoform switching during cell differentiation. Nat Biotechnol. 2010;28 (5):511-5. doi: 10.1038/nbt.1621. PubMed PMID: 20436464; PubMed Central PMCID: PMC3146043.

49. Trapnell C, Roberts A, Goff L, Pertea G, Kim D, Kelley DR, et al. Differential gene and transcript expression analysis of RNA-seq experiments with TopHat and Cufflinks. Nat Protoc. 2012;7 (3):562-78. doi: 10.1038/nprot.2012.016. PubMed PMID: 22383036; PubMed Central PMCID: PMC3334321.

